# Do working forest easements work for conservation?

**DOI:** 10.1101/2023.08.24.554638

**Authors:** Jonathan R. Thompson, Alexey Kalinin, Lucy G. Lee, Valerie J. Pasquarella, Joshua Plisinski, Katharine R. E. Sims

**Affiliations:** Harvard Forest, Harvard University; Economics Department, Environmental Studies Department; Amherst College

**Keywords:** Conservation Easement, Forestry, Land Protection, Land use, Private Land

## Abstract

Conservation easements are voluntary legal agreements designed to achieve conservation goals on private land. Extensive public and private funding has been used to establish “working forest” conservation easements (WFCE) that aim to protect conservation values while maintaining commercial timber production. We use variation in the timing and location of easements to estimate the impacts of WFCEs in Maine from a 33-year time-series of forest loss and harvesting. Compared to matched controls, WFCEs reduced forest loss rates by just 0.0004% yr^-1^, the equivalent of 3ha yr^-1^, and increased the rate of harvesting by 0.37% yr^-1^, or 2935ha yr^-1^ within the 839,000 ha enrolled. More recent easements contained stricter restrictions on harvest practices and these easements reduced harvest by 0.62% yr^-1^. Overall, WFCEs supported continued harvests, but did not appear to provide substantial ecological benefits. Future easements could be more effective if they include additional provisions for harvest restrictions and monitoring.

## 1.0 Introduction

Protection of private land is increasingly needed to achieve societal goals for climate regulation and maintenance of biodiversity, ecosystem functions, and natural resources. Conservation easements are a primary mechanism for advancing public conservation goals on private land (Parker & Thurman, 2019). Easements transfer a subset of property rights from the landowner to a land trust or government agency and restrict specified land uses in perpetuity (Rissman et al., 2013). Easements are often preferred for conservation because they are less expensive than fee acquisition and the landowner retains most property rights while receiving substantial income and/or favorable tax treatment.

The proliferation of conservation easements in the U.S. began in the early 1980s when Congress incentivized their use through changes to the tax code and passed the Uniform Conservation Easement Act (USCA) (Kay, 2016). The U.S. federal government now spends more than $450-million annually to conserve agricultural and forest lands using conservation easements (CRS, 2017; CRS, 2018) and grants more than $1-billion in tax breaks for new easements (CRS, 2019). Today, more than 16-million hectares are protected by conservation easements—more than the combined area of all U.S. National Parks. Despite their vast extent, few studies have examined the impact of easements on conservation outcomes or land-use practices (but see Nolte et al., 2019).

Specific land-use restrictions within conservation easements are individually negotiated between the landowner and the easement holder but must achieve at least one of the broadly defined conservation purposes articulated by the Internal Revenue Service and the USCA. Eligible purposes include outdoor recreation, fish and wildlife habitat, open space scenery, and historic preservation. So-called “working forest” easements are an informally defined class of easements intended to protect managed forests from conversion to non-forest uses and/or to promote sustainable forestry and other conservation values (Tesini, 2009). In 1990, the Forest Legacy Program was added to the U.S. Farm Bill to help fund the protection of privately-owned working forests through conservation easements and land purchases (Murray et al., 2018). A core objective of the Program, and of working forest easements generally, is to protect timber-based economies and the rural culture and landscapes they support, while also enhancing ecological conditions and recreation opportunities by protecting long-term access and large contiguous land parcels (Noone et al., 2012; Legaard et al., 2015; L’Roe & Rissman, 2017; Reeves et al., 2020). Since the establishment of the Forest Legacy Program, the largest share of forestland protected in the U.S. has been through WFCEs.

WFCEs vary substantially in terms of the protections they provide. Most focus on limiting parcelization and ensuring continued forest cover, and Forest Legacy Funded easements require multi-resource management plans (USFS 2017). Some easements include restrictions meant to curb unsustainable harvesting, including temporary harvest moratoriums, limits on timber volume removed relative to growth, and protections in sensitive areas (Owley & Rissman, 2016; Daigle et al., 2012). Proponents argue that WFCEs can contribute both to conservation objectives and to local and regional economic prosperity (Murray et al., 2018). Opponents argue that the protections offered by these easements offer little public benefit because they do not address unsustainable harvest practices and are often applied where threats of conversion are low (Pidot, 2005). The harshest opponents brand them as public subsidies to industry that primarily benefit distant investors (Sayen, 2023).

The proliferation of large WFCEs in the late 1990s coincided with the broad scale financialization of timberlands. Land financialization is the process of incorporating land into global networks of investment capital as financial assets (Gunnoe et al., 2018). Today, nearly all vertically structured timber and wood products commercial timberland in the U.S. have been acquired by investment interests, including timber and real estate investment trusts (Bliss et al., 2010; L’Roe & Rissman, 2017; Gunnoe et al., 2018).

Concerns that the shorter time horizons of investment owners might incentivize them to break-up, convert, and/or mis-manage the forests have led many conservationists to support establishment of large WFCEs (e.g., Stein, 2011). WFCE were also attractive to the new fiduciary owners, who could enhance short-term returns from the land investment from selling the restrictions on development (Saul, 2021).

To assess the impacts of WFCEs on land-use change and management practices, we use a 33-year time series of forest loss and harvesting data in the state of Maine (Figure 1). Maine serves as an ideal case study, having experienced rapid growth in land conserved through WFCEs during this period. Indeed, in 1990, just six percent of the Maine landscape had any type of protection and 80-percent of that was public land. By 2020, 22-percent of Maine had a protected status with large WFCEs accounting for 70-percent or 839,000 ha, approximately the size of Yellowstone National Park.

**Figure 1.**
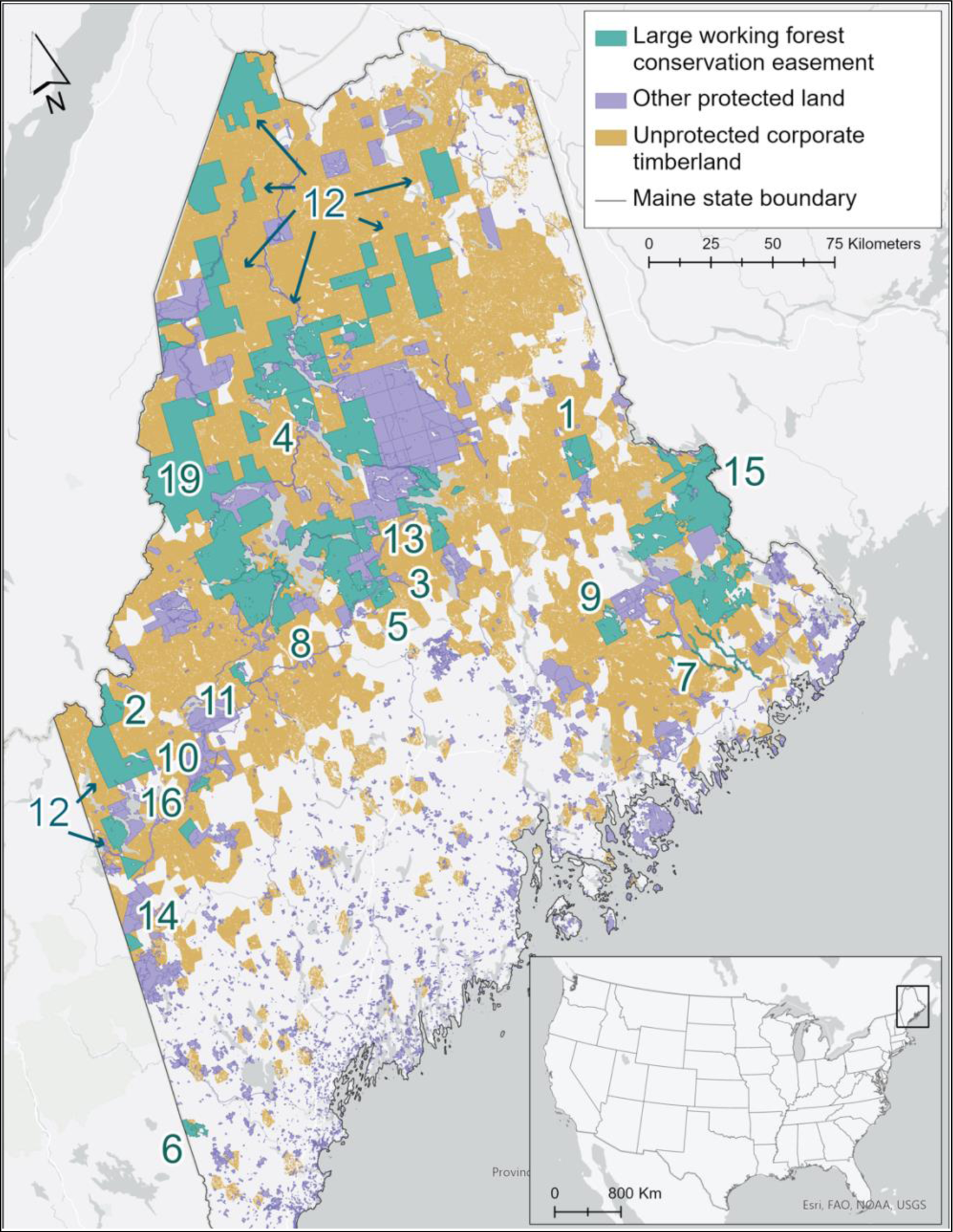
We analyzed rates of forest loss and harvest within 19 large working forest conservation easements (green) that span 839,000 hectares and are embedded in a matrix of other protected (lavender) and unprotected corporate owned timberlands (brown) throughout northern Maine. Note the numbers on the map correspond to the WFCEs described in Table 1.

**Table 1.** Attributes of the 19 working forest conservation easements analyzed in this study. Note the row numbers correspond to the numbers on the map inFigure 1.

| # | Easement Name | Year | Area (ha) | Fee owner/Grantor | Easement Holder | Holder Type |
| --- | --- | --- | --- | --- | --- | --- |
| 1 | Apple Conservation | 2016 | 12,985 | Shelterwood Holdings, I, LLC | Forest Society of Maine | NGO |
| 2 | Boundary Headwaters | 2004 | 9,105 | SP Forests, LLC; International Paper | Forest Society of Maine | NGO |
| 3 | Gulf Hagas - White Cap | 2016 | 2,855 | Pine State Timberlands, LLC | Maine Bureau of Parks & Lands | State |
| 4 | Katahdin Forest Project | 2002 | 75,808 | Maine Timberlands Company | The Nature Conservancy | NGO |
| 5 | Katahdin Ironworks | 2009 | 10,828 | AMC Maine Woods, Inc. | Maine Bureau of Parks & Lands | State |
| 6 | Leavitt Plantation Forest | 2003 | 3,405 | GMO Forestry Fund 1, LP | Maine Bureau of Parks and Lands | State |
| 7 | Machias River Watershed | 2003 | 5,348 | SP Forests, LLC; International Paper | Maine Atlantic Salmon Commission | State |
| 8 | Moosehead Lake | 2012 | 143,402 | Plum Creek Maine Timberlands, LLC | Forest Society of Maine | NGO |
| 9 | Nicatous Lake | 2000 | 8,725 | Robbins Lumber, Inc. | Maine Bureau of Parks & Lands | State |
| 10 | Orbeton Stream | 2014 | 2,318 | Linkletter Timberlands LLC | Maine Bureau of Parks & Lands | State |
| 11 | Pierce Pond | 1996 | 2,722 | SD Warren Company | United States of America | Federal |
| 12 | Pingree | 2001 | 305,681 | Six Rivers Limited Partnership; Pingree Associates, Inc. | New England Forestry Foundation, Inc. | NGO |
| 13 | Roaches Pond | 2009 | 7,143 | AMC Maine Woods, Inc. | Maine Bureau of Parks & Lands | State |
| 14 | Robinson Peak Forest | 2008 | 2,720 | LBA Forest Stewardship Initiative, | Maine Bureau of Parks & Lands | State |
| 15 | Sunrise Easement | 2005 | 124,331 | Typhoon LLC | New England Forestry Foundation, Inc. | NGO |
| 16 | Tumbledown Mount Blue | 2002 | 3,148 | Hancock Land Company, Inc. | Maine Bureau of Parks & Lands | State |
| 17 | Upper St. John River North | 2012 | 1,694 | The Tall Timber Trust | The Nature Conservancy | NGO |
| 18 | Upper St. John River South | 2009 | 2,878 | Stetson Timberlands, Inc. | The Nature Conservancy | NGO |
| 19 | West Branch Penobscot Headwaters | 2003 | 114,046 | Merriweather, LLC | Forest Society of Maine | NGO |

We compare rates of forest loss and harvest within the easement areas to matched control sites in unprotected private lands that are similar in terms of relevant observable covariates. We also characterize and assess land-use restrictions articulated within the easements and how changes in easement strictness relate to observed patterns of land use.

Overall, we find that the easements had little effect on rates of forest loss or harvesting. Our analysis of easement restrictions indicates that more restrictions have been imposed over time, and that stricter easements reduced harvest frequency. These findings indicate that WFCEs have not produced substantial additional ecological benefits by reducing forest loss or harvest rates compared to status quo commercial timber operations. Effectiveness could be increased with changes in easement design and enforcement that prioritize areas under active threat from conversion and strengthen restrictions on unsustainable harvest practices.

## 2.0 Methods

### 1.1 Study-area and Easements

We analyzed WFCEs throughout Maine (Figure 1) where >7 million hectares of forests cover >90% of the land area. The southern half of Maine is dominated by northern hardwood forests, which transition to boreal spruce-fir forests in the north (Thompson et al., 2013). Two centuries of saw log harvesting and one century of intensive pulp production have created a forested landscape that is highly fragmented, traversed by roads, and dominated by even-aged trees (Gunn et al., 2019). Maine’s forests are primarily owned by private entities, including corporations (59%) and families (32%; Sass et al., 2021).

We define large WFCE as those protecting >1500 hectares of forestland and where maintaining forest management is one of the stated purposes of the easement. Using a geodatabase of all protected land in New England (Harvard Forest, 2023), and by obtaining easement contracts from the Maine Registry of Deeds, we identified 19 easements that met our definition of WFCEs (Table 1). These range in size from 1700 to > 308,000 ha and total 839,000 ha. They represent 49% of the total protected areas in the state, and 72% of the private protected area. Nine of the easements are held by conservation NGOs and ten by government agencies. At least twelve of the nineteen projects received public funding totaling > $50 million (Table 1), most of which (55%) came through the Forest Legacy Fund (Table S2).

### 1.2 Quantifying Land-use and Land-Cover change

Our panel dataset spans 1985 to 2018 and measures rates of forest loss and harvesting within the WFCEs and similar unprotected control sites. Landowner data came from Sass et al., (2020). Our unit of analysis is a 10-hectare hexagonal grid cell. We calculated the proportion of forest loss and forest harvest within each unit and year.

Our estimates of forest loss are based on the U.S. Geological Survey’s Land Change Monitoring, Assessment, and Projection (LCMAP; Brown et al., 2020). LCMAP is a Landsat-based (30m resolution) annual time series of land-cover produced using the Continuous Change Detection and Classification approach (Zhu & Woodcock, 2014). We used LCMAP CONUS v1.1 and re-classified pixels that transitioned from forest or shrub cover to a developed class as forest loss.

Our forest harvest estimates are based on annual maps of harvest events in Maine from 1985 to 2018 (Pasquarella et al. in review)^1^. Harvests were detected using the LandTrendr temporal segmentation algorithm (Kennedy et al., 2018) applied to Landsat time-series of three spectral indices: Normalized Burn Ratio, Normalized Difference Moisture Index and Tasseled Cap Wetness time series. The three sets of segmentation results were ensembled using a simple decision tree approach, which resulted in an accurate characterization of harvest events. See Supplemental Methods for details.

### 1.3 Environmental Variables and Covariate Pre-matching

To evaluate impacts of easements, we identified social and environmental variables known to be correlated with rates of forest loss and harvesting rates (based on Thompson et al., (2017a, 2020)) and used covariate pre-matching to ensure that the joint distributions of these variables were similar between treatment (WFCEs) and control groups. Matching variables for harvest rates include: correlates with forest productivity (i.e., light, climate, forest type, forest structure, nitrogen deposition), and with harvest access (i.e., topography, road density, wetlands, streams). Matching variables for forest loss include many of these same variables, plus distance to population centers and distance to energy transmission infrastructure (Table S3). Matching of hexagonal grid cells was done separately for each easement (Ho et al., 2011).

Matched hexagons for the harvest analysis were selected from the population of unprotected commercially owned timberlands, while matched hexagons for the forest loss analysis were selected from the population of all unprotected private land within the counties containing WFCEs. For each easement, matching was done without replacement, with similarity defined by nearest-neighbor measured in Malahanobis distance, with exact matching used for categorical variables (i.e., forest type) and calipers set to one standard deviation for continuous variables. Matching succeeding in balancing covariate distributions (Figure S4, S5). Trends in loss and harvest are similar prior to easement adoption dates as expected for well-matched sets (Figure 2, 3, Figure S2, S3).

**Figure 2.**
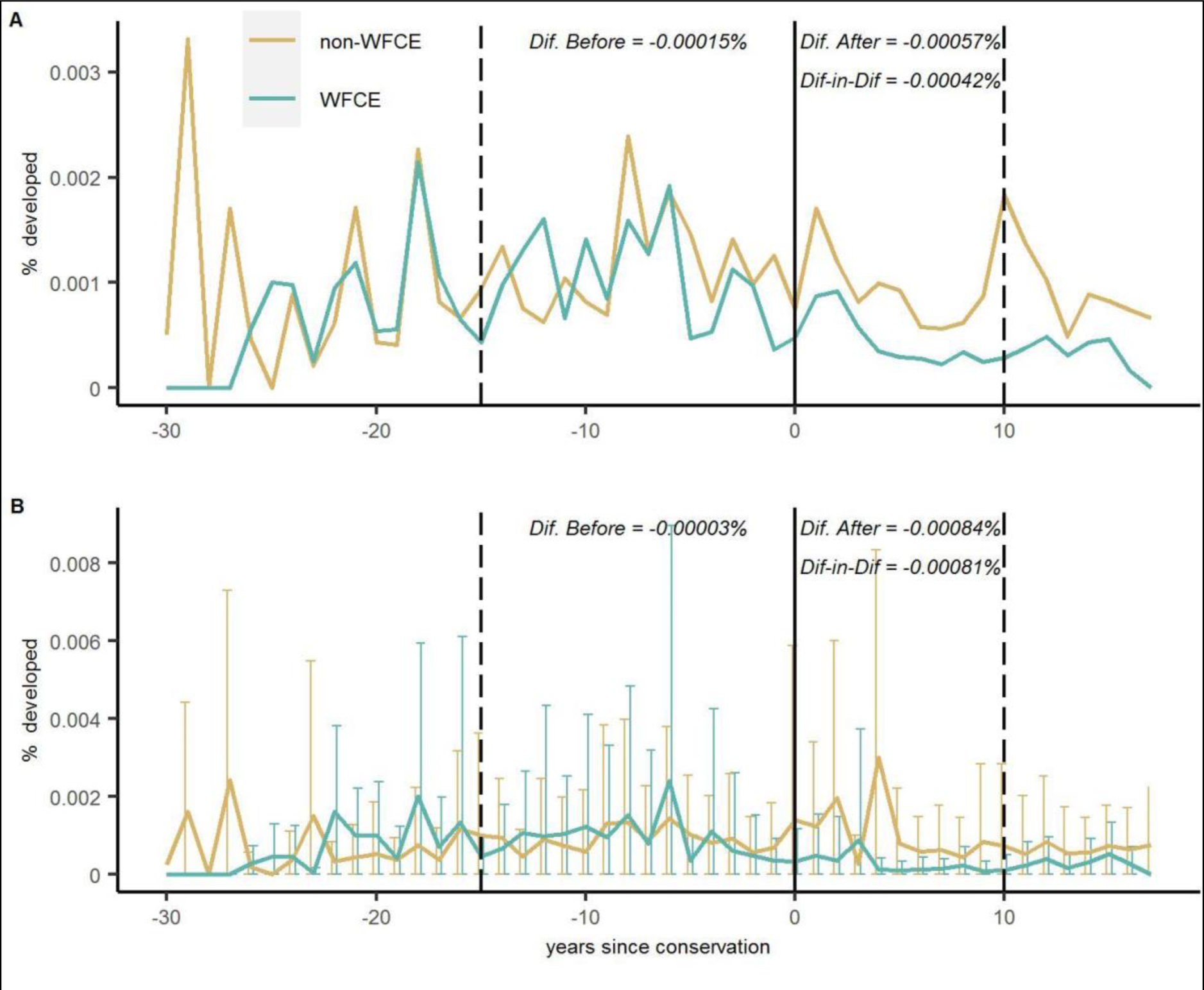
Annual rate forest loss within 19 working forest conservation easements (WFCE; blue) and their matched controls (brown). Vertical line indicates the year of conservation, which has been scaled to zero for all easements. **(Top)** Annual percent of forest loss calculated across all WFCE land, which effectively weights the easements by their size. **(Bottom)** Average annual percent forest loss within each easement, such that each easement is one sample regardless of size. Error bars are standard errors. The dotted lines at −15 and +10 indicate the years used to calculate the difference in difference estimates. The longer window of time for the pre-period reflects greater data availability for that time so we can reasonably extend the averaging window.

### 1.4 Estimates of Easement Impacts

We estimate the impact of the easements using a difference-in-differences approach, wherein we estimate the difference in forest loss and harvesting between the control and easement sites before easements were established and subtract that from the difference after the easements were established (also Figures 2 and 3). This approach accounts for general trends in conversion or harvest, so the estimated effects can be attributed to the easement policy.

**Figure 3.**
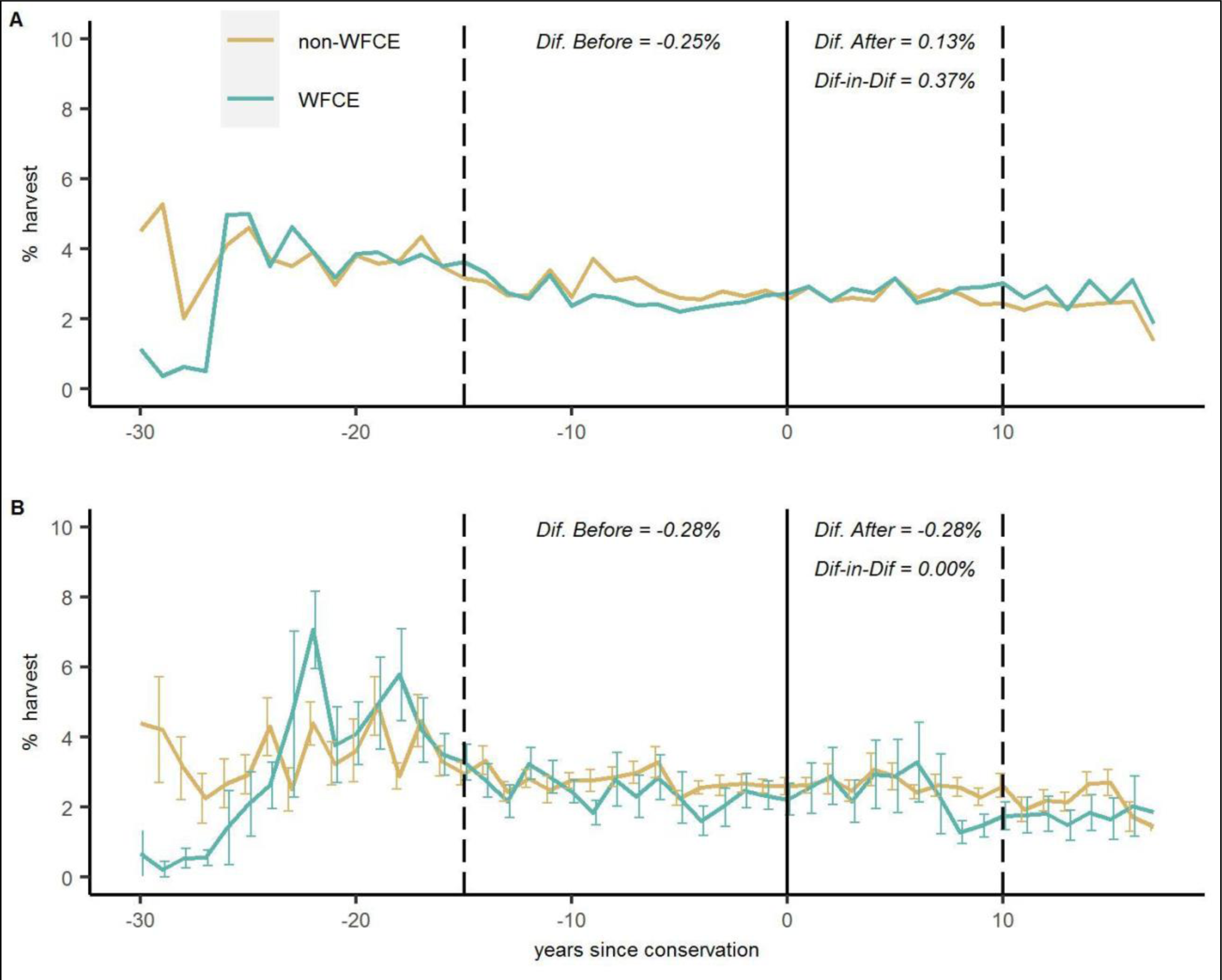
Annual rate of forest harvesting within 19 working forest conservation easements (WFCE; blue) and their matched controls (brown). Vertical line indicates the year of conservation, which has been scaled to zero for all easements. **(Top)** Annual percent of forest harvested calculated across all WFCE land, which effectively weights the easements by their size **(Bottom)** Average annual percent forest harvest within each easement, such that each easement is one sample regardless of size. Error bars are standard errors. The dotted lines at −15 and +10 indicate the years used to calculate the difference in difference estimates.

We evaluated these impacts with two sets of weights. First, we calculated the percent of WFCE area affected by each land use across all the areas within WFCEs, which weights the easements by their size and shows their aggregate effect within the region. Second, we evaluated the percent of forest loss at the individual WFCE scale, which gives each easement equal weighting as a treatment unit. We also compare rates of loss in WFCEs to overall rates in the state as a whole (Figure S1).

### 1.5 Easement Coding

Following Owley and Rissman (2016), who examined trends in easement complexity by coding easement terms according to the degree of control, we coded terms in the 19 WFCE contracts related to easement purposes, allowed land uses (e.g., forestry practices & construction), and management, enforcement, or conservation outcomes (e.g., subdivision & subdivision cost, ecological monitoring). To construct an index of strictness, we used the 14 terms that captured potential restrictions on land use or harvesting. Each term has possible values ranging from 0 to 1, with language that corresponds to more control over landowner activity receiving higher scores; these are summed to create the “easement strictness index.”

## 2.0 Results

Compared to Maine’s forests as a whole, corporate-owned timberlands, whether protected by a WFCE or not, have a low rate of conversion to non-forest land cover (Figure S1). Using the difference-in-difference approach to compare rates of forest loss between easement and matched control units at the WFCE scale, we find a very small reduction—i.e., 0.0008% less forest loss per year (Figure 2). When weighting by the area of the easements, the reduction is even smaller—i.e., 0.0004% less forest loss per year. Assuming the control sites offer a reasonable counterfactual and scaling by the areas of the easements, we estimate that the WFCEs prevented 3.17 ha yr^-1^ from being converted. No individual WFCE clearly altered the trajectory of forest conversion relative to their control plots (Figure S2).

Unlike forest loss, rates of forest harvesting within WFCEs and in unprotected corporate forestlands are higher than for Maine’s privately-owned forests overall (Figure S1). There is substantial variation in harvest rates across the WFCEs, and few show distinct changes in harvest rates after the time of easement (Figure S3). The Roach Pond Track is one exception; this property was bought and eased by a conservation organization (AMC) with the explicit intention of letting it recover from overharvesting before resuming harvesting in the future, which is reflected in the lower rates of harvest after the easement date (Figure S3). Overall, comparing rates of forest harvesting between easement and control units with equal weighting of easements, we estimate that easements resulted in no change in harvest rates—i.e., a difference in difference estimate of 0.00% harvested area per year (Figure 2). When weighting by the area of the easements, we estimate WFCEs increased harvest rates by 0.37% per year. Using the areas of the easements, we estimate the WFCEs resulted in 2935 ha yr^-1^ more harvesting.

There is substantial variation in easement language concerning restrictions on land-use, monitoring, and harvesting practices (Figure 4, SI methods). We find a general trend towards stricter easements over time (Figure 4) with additional provisions in easement language for reporting, monitoring, and controls on harvest practices. We also find that harvest rates declined relative to their matched controls after adoption of restrictive easements (highest 50^th^-percentile of strictness index) by 0.62% yr^-1^ (Figure 4). Harvest rates within WFCEs with low strictness (lowest 50^th^-percentile) increased by 0.61% yr^-1^.

**Figure 4.**
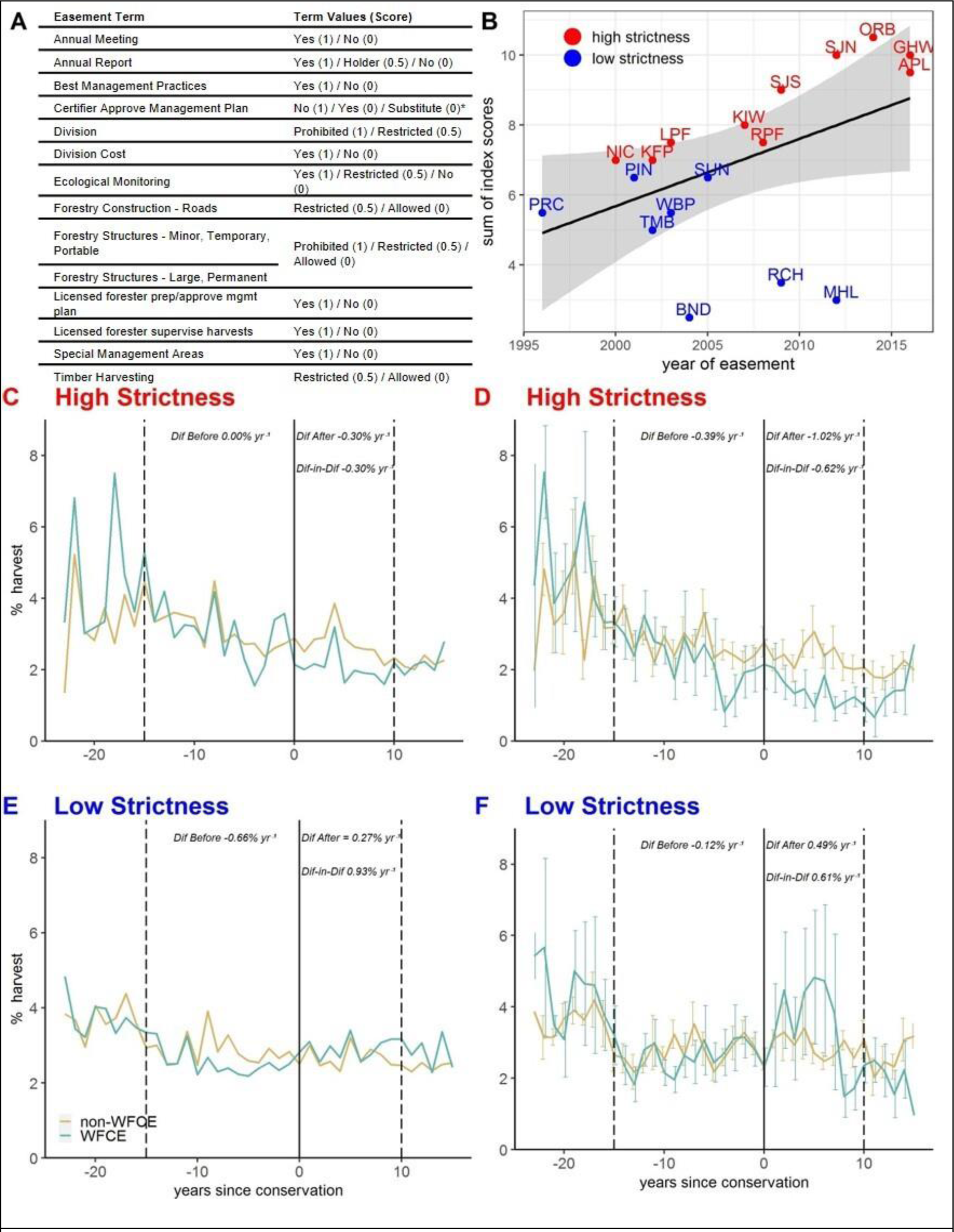
A. Coding rubric for harvesting restrictions used to make the strictness index. **B.** Harvest strictness index over time. **C.** Harvest rates within all easement land within the upper 50^th^ percentile of easement strictness and matched controls **D.** Harvest rates measured per easement for the easements in the upper 50^th^ percentile of easement strictness and matched controls (error bars = standard errors) **E.** Harvest rates within all easement land within the lower 50^th^ percentile of easement strictness and matched controls **F.** Harvest rates measured per easement for the easements in the lower 50^th^ percentile of easement strictness (error bars = standard errors) and matched controls.

## 3.0 Discussion & Conclusion

A primary goal of WFCEs has been to prohibit permanent development and ensure future forest cover. While we find that WFCEs have experienced low rates of conversion, they are only slightly lower than the counterfactual sites within unprotected private land generally (∼ 3 ha yr^-1^). This is consistent with the fact that the lands protected by WFCEs are overwhelmingly located in the rural unincorporated territories of Maine, where there are few people and there has been a low threat of conversion to date. Near-term benefits of WFCEs might be improved by prioritizing new easements on lands that are at the highest threat of conversion.

Economic and ecological pressures are shifting with time and the threat of forest loss may increase in the future. Like all easements, the terms of these WFCEs are attached to the land in perpetuity, including through ownership transitions. Over the past thirty years, the major driver of forest loss in the region has been dispersed residential construction associated with exurban development (Thompson et al., 2017b). In contrast, the limited forest loss we did observe in unprotected lands arose from the development of energy infrastructure, e.g., powerlines, pipelines, wind, and solar. Given that current plans to decarbonize and electrify energy sectors are expected to dramatically alter landscapes worldwide (Lovering et al., 2022) and in Maine specifically (Merrow, 2018), greater impacts of the WFCEs may also be realized during future climate adaptation and mitigation efforts.

WFCEs also have the potential to improve forest management practices to benefit wildlife habitat and ecosystem services (L’Roe & Rissman, 2017). However, we find that the establishment of the WFCEs overall had little impact on the rate of harvesting. This is consistent with the stated goals of many easements to maintain harvesting, but disappointing given that rates of harvest in the region are ecologically degrading the forests (e.g., Gunn et al., 2019). We do find that the restrictions of harvest practices within the WFCEs have increased over time and that stricter easements reduced the rate of harvesting. However, because the stricter easements tend to be the most recent, there is substantial uncertainty in these estimates.

Nonetheless, these new WFCEs demonstrate potential approaches to target threats from unsustainable and/or damaging harvesting. For example, the 2016 Apple Easement in Reed Plantation, Maine, includes explicit restrictions to maintain sustainable harvest volumes and practices and includes protections on water quality and fish and wildlife habitat as well as endowed funds for easement monitoring. It is possible that WFCEs meaningful change forestry practices, even if they don’t alter the rate of harvesting. There is some evidence to support this—e.g., Zhao et al., (2020) report larger average harvested tree size within private protected land in Maine (but not explicitly WFCEs), suggesting landowners may have longer harvest rotation periods in protected forests. In addition, WFCEs may have contributed other economic benefits through recreational and heritage use of these lands (Murray et al. 2018, Daigle et al. 2012).

How should conservationists interpret these findings? As scarce public conservation dollars are used to buy WFCEs from financial firms and conservation organizations tout the land as “protected”, many might reasonably assume that the forests are protected from the land uses that threaten them most. Yet, our results suggest a misalignment between the application of funding and restrictions in the easements and the current nature of the threats to these forests. The threat of forest conversion in the region is extremely low but is the primary focus of the WFCEs. Forest harvesting practices are well-documented threats and yet the easements include few restrictions on harvesting. We also do not find evidence that easements were needed to sustain forestry in the region; the unprotected forests have similar rates of harvest.

However, we do find that newer WFCEs are more likely to restrict unsustainable and damaging harvest practices. Threats to forests are also changing over time and conversion pressure may be higher in the future. Overall, we conclude that private philanthropy and public funding like the Forest Legacy Program—whether in Maine or elsewhere--could increase conservation benefits by requiring WFCEs to better align easement protections with the land-use practices that most threaten those lands.

## Acknowledgements

This work was supported by USDA/NIFA Grant No. #2021-67023-34491, NSF Grant No. DEB-LTER-18-32210, NSF-DISES Grant 22-05705 and a Lone Mountain Fellowship from the Property and Environment Research Center to the lead author. We thank Colin Brown of Downeast Lakes Land Trust for valuable insight into the conservation landscape of the Downeast Lakes region. We also thank Kirston Buczak of the Forest Legacy Program and Sarah Demers and Liz Petruska of the Land for Maine’s Future Program for providing data on their respective programs. L. Mapes, D.R. Foster, J. Daigle, R. Perschel, J. Leahy, and A. Daigneault provided valuable feedback on drafts of this manuscript.

## Supplementary Materials

### Methods

#### Harvest mapping

We used the Google Earth Engine implementation of LandTrendr temporal segmentation approach (Kennedy et al., 2018, 2010) to generate inputs for our harvest event detection ensemble. We applied LandTrendr to annual medoid composites of all high-quality Landsat 5, 7 and 8 Collection 1 Surface Reflectance observations acquired between June 20 and September 20 (Northern Hemisphere growing season) for the years 1985–2020 using the parameters shown in Table S1. Initial segmentation results were generated separately for three SWIR-based indices, (1) the Normalized Burn Ratio (NBR), (2) the Normalized Difference Moisture Index (NDMI), and (3) Tasseled Cap Wetness (TCW). LandTrendr outputs include a series of segments, which correspond to relatively stable periods, and vertices, which were identified as inflection points along a spectral trajectory and are indicative of potential changes in surface conditions (Pasquarella et al., 2022). We considered all loss vertices, i.e., those with spectral changes in the direction of decreased vegetation cover, as potential disturbance events. To differentiate harvests from longer-duration disturbances such as those related to drought or forest insect damage, we removed vertices associated with segments greater than two years in duration, leaving only short-term (<2 year) events that are more likely associated with harvesting.

We used all available FIA field plot measurements collected in the state of Maine between 1999 and 2019 to train and cross-validate harvest detection ensembles and we had access to true plot locations through a memorandum of understanding between the USFS and Harvard University (MOU #09MU11242305123). Our Maine FIA dataset consisted of 13,299 measurements (i.e., unique space-time coordinates) recorded for 3,265 plots (i.e., unique spatial locations), and of these, we analyzed the 3,220 FIA plots that had been remeasured at least once and our final dataset included 10,034 pairs of sequential FIA measurements, of which 1,711 recorded basal area removal (harvest).

To integrate the Landsat-based and FIA datasets, we queried the LandTrendr results for all years between the first and second FIA measurement years to determine if a potential harvest event was detected between measurements. If multiple events (vertices) were detected in a given measurement period, we recorded information for the event with the greatest magnitude of change. We excluded the first year of each measurement pair to prevent double-counting endpoint years for plots with two or more remeasurements. The resulting dataset included a record for each FIA remeasurement with plot information from each measurement pair (m_a_ and m_b_) as well as the LandTrendr features for each of the three spectral indices we considered.

We used a degenerate decision trees approach to combine LandTrendr results for different spectral indices into a final time series of forest harvest event maps. Degenerate trees are a subclass of binary trees where each decision node has only a single parent node. We provide an example of our DDT implementation at <u>github.com/valpasq/lt-ensemble</u>, including a Python notebook with example functions for running a sweep over series of thresholds and determining optimal thresholds for each feature as well as an Earth Engine script for applying thresholds to LandTrendr results (Pasquarella, 2022).

We used our FIA remeasurement dataset to perform a three-fold cross validation replicated ten times for a total of 30 folds. We found that we were able to detect harvest events that removed at least 30% of total basal area with an F1 score of 0.72 (σ = 0.02) with a false negative error rate (omission) of 0.32 (σ = 0.02) and false positive error rate (commission) of 0.23 (σ = 0.03). Visual comparisons of our mapped suggest similar spatial patterns to v2020-5 forest disturbance products produced as part of the U.S. Forest Service’s Landscape Change Monitoring System (LCMS) project (USDA Forest Service, 2021), which was also considered as a potential data source for this study. However, the LCMS dataset does not distinguish among forest harvest and other “fast loss” events such as harvest, mechanical, wind/ice, hydrology, and debris, and was found to have notably lower F1 scores (0.62), with a relatively low false positive rate (0.13), but a very high false negative rate (0.54) when compared to the same reference points. Although it might be assumed that national LCMS products would be better suited for detecting higher-intensity stand-replacing disturbances, higher rates of omission were observed across all percent basal area removal categories in the FIA-based analysis, suggesting that using fast loss classifications from the LCMS dataset to represent potential harvest events would have resulted in underestimation of harvest rates with important implications for the assessments in this study. Thus, improvements in harvest event detection accuracy justified the development of custom ensemble models for this project rather than relying on existing datasets as we did for evaluating forest conversion/loss.

#### Forest Loss Mapping

We incorporated Landover change data from the United States Geological Services (USGS) Land Change Monitoring Assessment and Projection (LCMAP) program. We used Conterminous Collection 1.2 from 1985-2019 as downloaded from https://eros.usgs.gov/lcmap/apps/data-downloads. This 30m resolution land cover classification product is derived from the Change Detection and Classification algorithm (CCDC) (Zhu and Woodcock 2014). CCDC uses all clear Landsat Analysis Ready Data (ARD) observations to fit harmonic regression models to predict surface reflectance. Mapped land cover changes are based on deviations from predicted reflectance values. We used the LCMAP’s Primary Land Cover (LCPRI) product, which represents the most likely land cover class as of July 1^st^ of each year. LCPRI classes include Developed, Cropland, Grass/Shrub, Tree Cover, Water, Wetland, Ice/Snow, and Barren. In this study, we calculated the area of “Tree Cover” that changed to anthropogenic land covers: “Developed” and “Cropland”. We converted all land cover and land cover change values into percent of the sample unit to account for non-uniform size.

#### Identifying Large Working Forest Conservation Easements

We identified large working forest conservation easements (WFCEs) in Maine using multiple sources of information in an iterative process: 1) protected open space layers from the Harvard Forest and others, 2) land ownership data from Sass et al. (2020), 3) project lists from federal and state conservation funding programs, 4) easement contracts, and 5) personal communication with easement holders.

WFCEs are identified within the New England protected open space (NE POS) layer (Harvard Forest, 2023). Prior to analysis, this layer was updated to include new data (circa May 2021) from state-provided conservation and open space layers as well as data from Loeb and D’Amato (2019). In any update of NE POS, the new data has a protected area (PA) “type” assigned to it based on 1) year of protection (protected 1990 or later), 2) ownership (public or private), 3) GAP status, and 4) size. LWFCEs belong to a “type” of PA that is defined by 1) year of protection 1990 or later, 2) private (for-profit) ownership, 3) GAP status of 3 or 4 (which indicates extraction is allowed), and 4) a size larger than 1,400 hectares. Both fee and easement PAs may meet this criterion; we focus on easement-protected PAs for this analysis due to the written record of allowed uses an easement provides. Given the small number of LWFCEs that meet this criterion, we confirm each WFCE individually.

To confirm PAs categorized as WFCEs were correctly classified, we compared our spatial and tabular data to the sources listed in paragraph one. Each LWFCE is unique, and the particular sources used to confirm WFCE status varied according to need and context. We compared the footprint and attributes for the WFCEs to other conservation layers including Maine’s Conserved Lands dataset and The Nature Conservancy’s Secured Areas. Project lists provided by the Forest Legacy Program and Land for Maine’s Future, which contain tract-level details on dates, locations, and sizes, were also used to confirm WFCEs. In cases where private ownership was certain but for-profit status was not, we used the Sass et al. (2020) ownership layers to provide additional context. In some cases, we had direct communication with easement holders to either confirm or retract WFCE status.

WFCEs that were categorized based on attributes in spatial data were most commonly removed from the WFCE group based on private non-profit ownership or management that is not focused on forestry. Community forests represent one type of PA that is commonly categorized as WFCE based on attributes in spatial data. Community forests are often large enough to be WFCE based on size, may be privately owned, and harvest trees for community benefit, thus having GAP 3 or 4. However, the motivation and management of community forests is completely different from a corporate-owned timberland (pers. comm.). Several tracts comprising the Downeast Lakes Community Forest were removed from the WFCE group based on this distinction following personal communication with a land trust representative.

To ensure that we did not miss any WFCEs, we investigated non-WFCE PAs in NE POS by querying the data for large (>1,400 ha), non-publicly owned PAs protected since 1990 with a GAP status greater than 2. PAs returned by this query were compared against the same information sources mentioned above. We also matched all the projects listed in the Forest Legacy Program and Land for Maine’s Future data that we did not have categorized as WFCE to other PAs to confirm they are not WFCE. No new WFCEs within the study period for this analysis were identified using these methods.

#### Identifying unprotected corporate lands

Unprotected, corporate-owned timberlands were identified using ownership data from Sewall (2018) and Sass et al. (2020) as well as HF POS (Harvard Forest, 2023). Sass et al. (2020) ownership data is gridded data of landowner types, rather than names, interpolated from Forest Inventory Analysis data. NE POS is used to erase protected areas from the ownership data.

#### Matching Variables

Table S2 provides details about the matching variables used for the forest loss and harvesting analysis. All data was rasterized (vector data) or resampled (raster data) to 30m and coregistered to a common spatial extent and projected coordinate system. Most of the variables were aggregated from 30m raster values to ∼10ha sample units by taking the mean of the ∼11 cells that fell within each hexigon. In some instances, it was more appropriate to calculate a percent or majority value (see Table S2).

#### Easement collection and coding

Easement collection happened in parallel to refining the list of WFCEs, as the easement contracts themselves contain information, sometimes including maps, that can be used to confirm WFCE status. Easements were obtained from the Maine Registry of Deeds and coded according to their terms. In total, we coded 48 pieces of information about each easement based on a coding scheme developed for general conservation easements (not necessarily working forest easements). This included basic information such as the year and holder type, the purposes of the easement, and the allowed uses. Easement coding was done in an iterative process, with some terms that relate specifically to working forests being coded in a second and third pass, as patterns in easement language revealed themselves. For example, in the first pass of reading easements we had only one column for coding the permissibility of forestry construction. Given the variation across WFCEs in forestry construction restrictions, this one column was not meaningful and was expanded into separate columns for road construction, road maintenance, minor/temporary/portable construction, and large/permanent construction. These more specific data points were created based on patterns in language across easements.

The possible values for each term depend on the nature of the term and are thoroughly documented in the associated metadata. Purposes are coded in binary presence/absence, while land uses may be “allowed,” “restricted,” “prohibited,” or “unknown” if easement language is inconclusive. Most of the terms we coded relate to land uses, including recreation, forestry, agriculture, and construction for those activities; rock, gravel, and water extraction; development for renewable energy, residential, commercial, or industrial purposes; and use of chemicals. We also coded terms related to (co-)management of the land, such as annual meetings and reports; division restrictions, costs, and withdrawn lots; third party certification; and the easement holder’s affirmative rights, including ecological and compliance monitoring.

In coding easement language, we find substantial variation in land-use restricts among WFCE. For example, while all easements require the easement holder be granted the general right of compliance monitoring, only 16% of the WFCEs specifically allow the easement holder to conduct ecological monitoring of the land. All WFCEs restrict subdivision to some degree, although most (79%) do ultimately allow subdivision. Approximately one in four WFCEs allow for residential development with restrictions. Over half (58%) have some restrictions on timber harvesting through designating special management areas, though these vary in definition and strictness; and 47% require harvest supervision by a licensed forester. Two-thirds of WFCEs (63%) prohibit large permanent forestry construction such as processing plants, though 32% allow such construction with restrictions, including three that allow lots to be withdrawn from the easement area to site such facilities. Most (79%) require an annual meeting between the landowner and the easement holder, while only 32% require the landowner to submit an annual operating report, and the vast majority (84%) restrict forestry construction to some degree.

A subset of 14 coded easement terms were converted to numeric values and summed to produce a “strictness index.” The 14 terms relate to management, division, harvest, and forestry construction and are described in Figure 4a. The terms were selected based on their relationship to and potential influence over harvest and conservation outcomes. We performed a simple linear regression of the strictness index scores with year of easement and found a significant relationship at 95% confidence.

---

**Figure S1.**
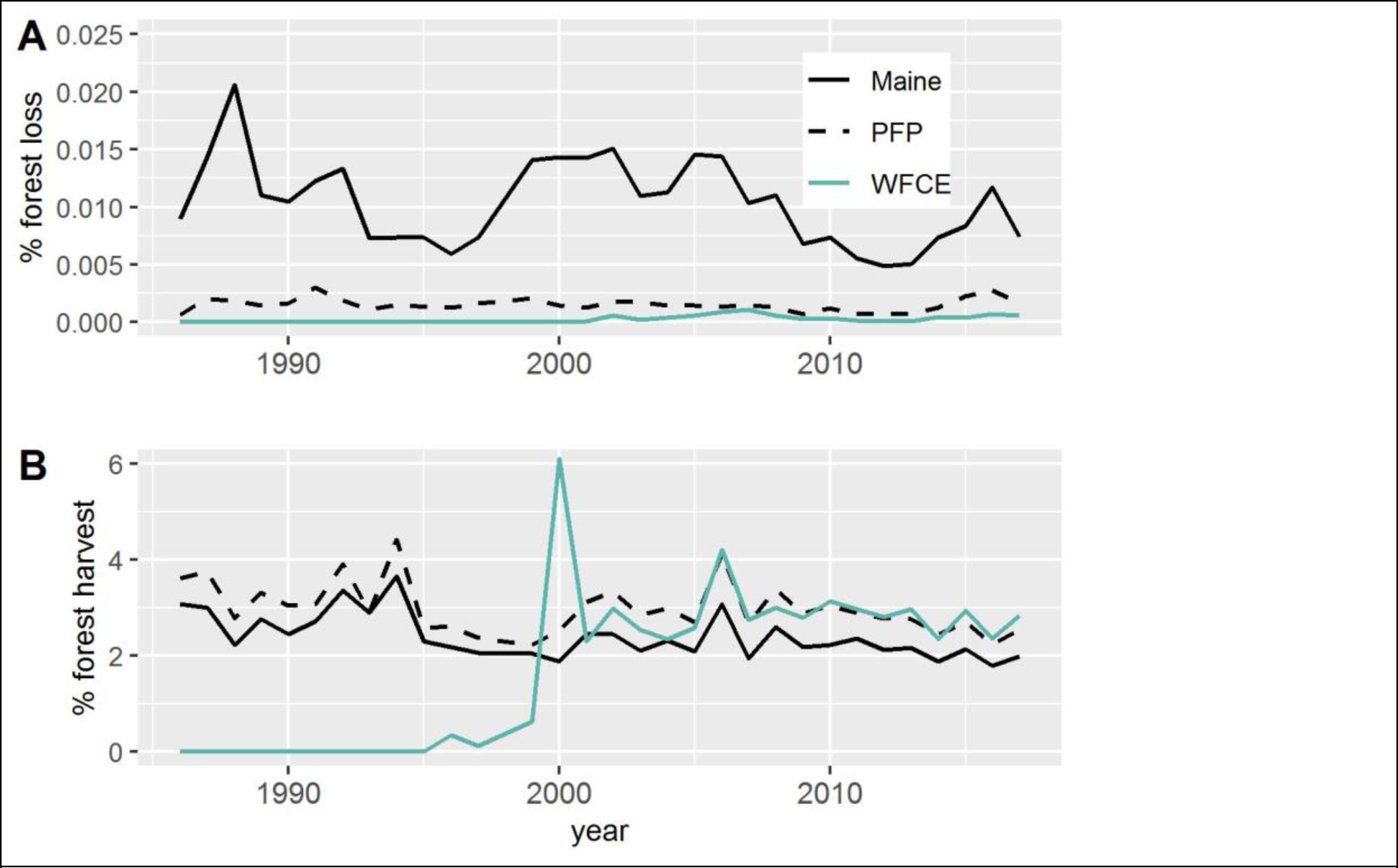
A) The annual percent of forest loss (A) and forest harvest (B) in all forests within the state of Maine (Black solid), the unprotected corporate-owned forests in Maine (Black dash), and the large working forest conservation easement (WFCE) in Maine (Green). The increase in WFCE harvests around the year 2000 coincides with the proliferation of new WFCE on the landscape.

**Figure S2.**
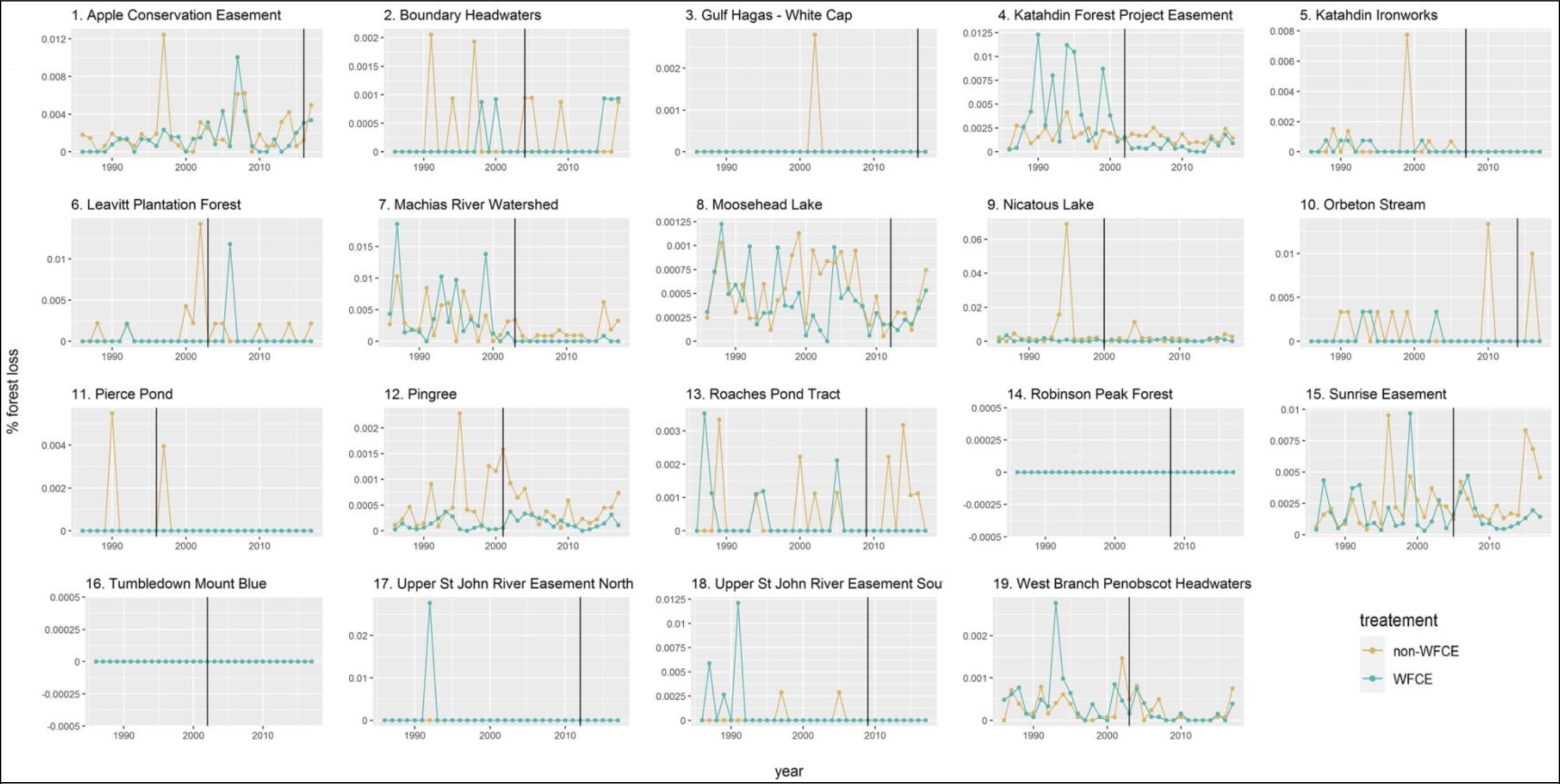
Rates of forest loss within 19 working forest easements (teal) and their matched controls (orange). The vertical line indicates the timing of the easement. In plots that appear to show only a line for WFCEs, the two lines are overlapping.

**Figure S3.**
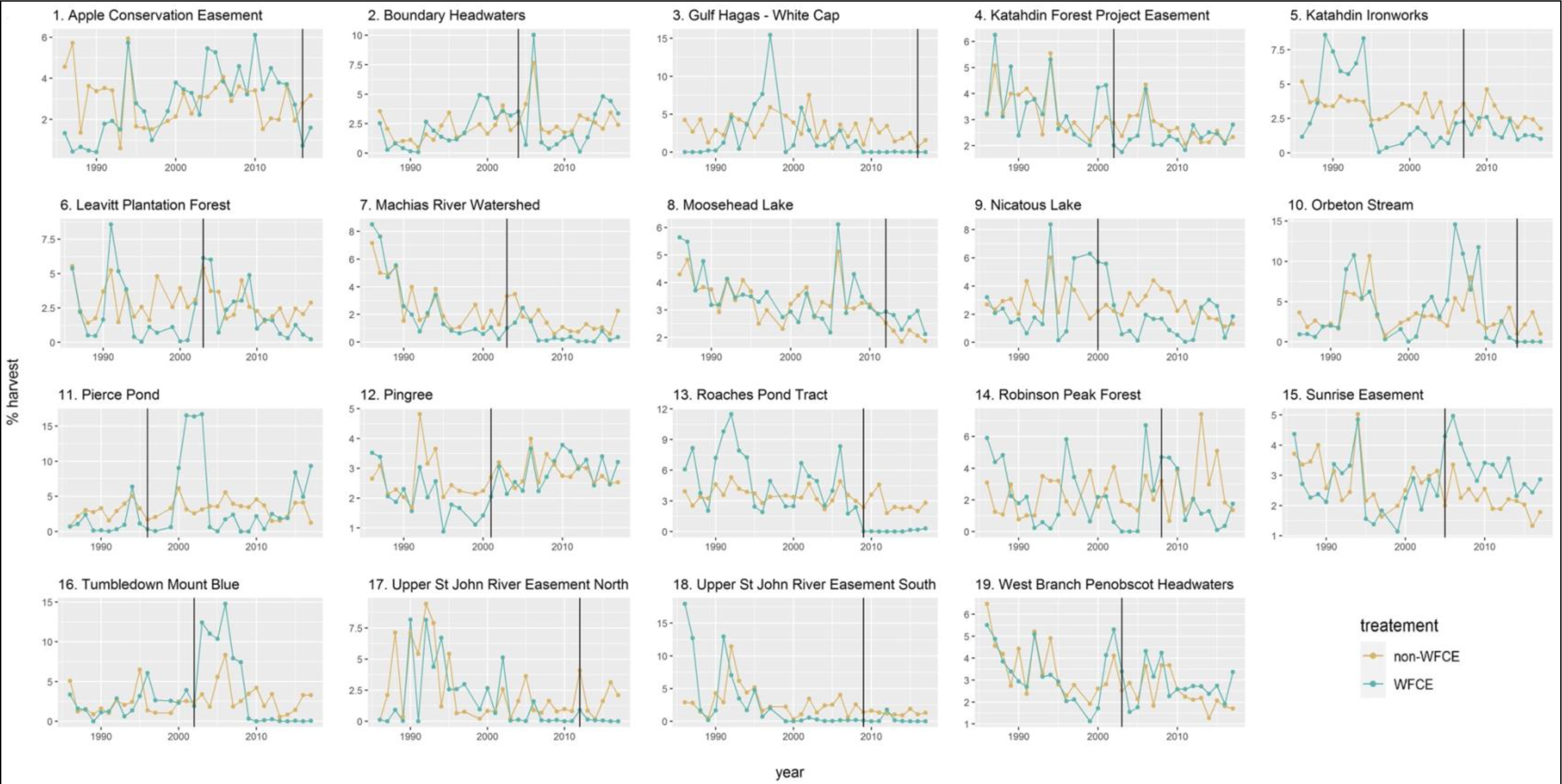
Rates of forest harvesting within 19 working forest easements (teal) and their matched controls (orange). The vertical line indicates the timing of the easement.

**Figure S4.**
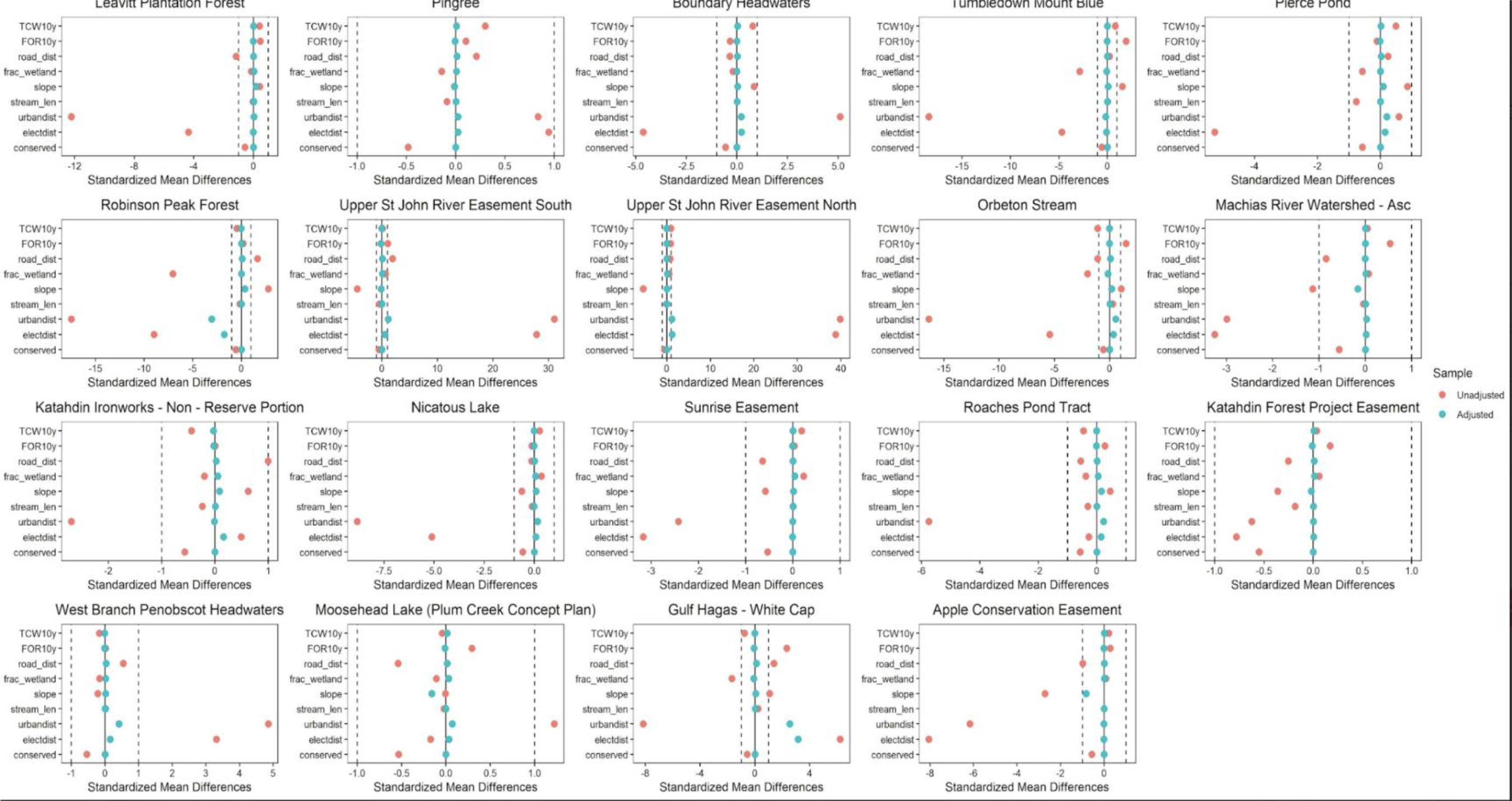
Balance plots for covariate pre-matching for forest loss.

**Figure S5.**
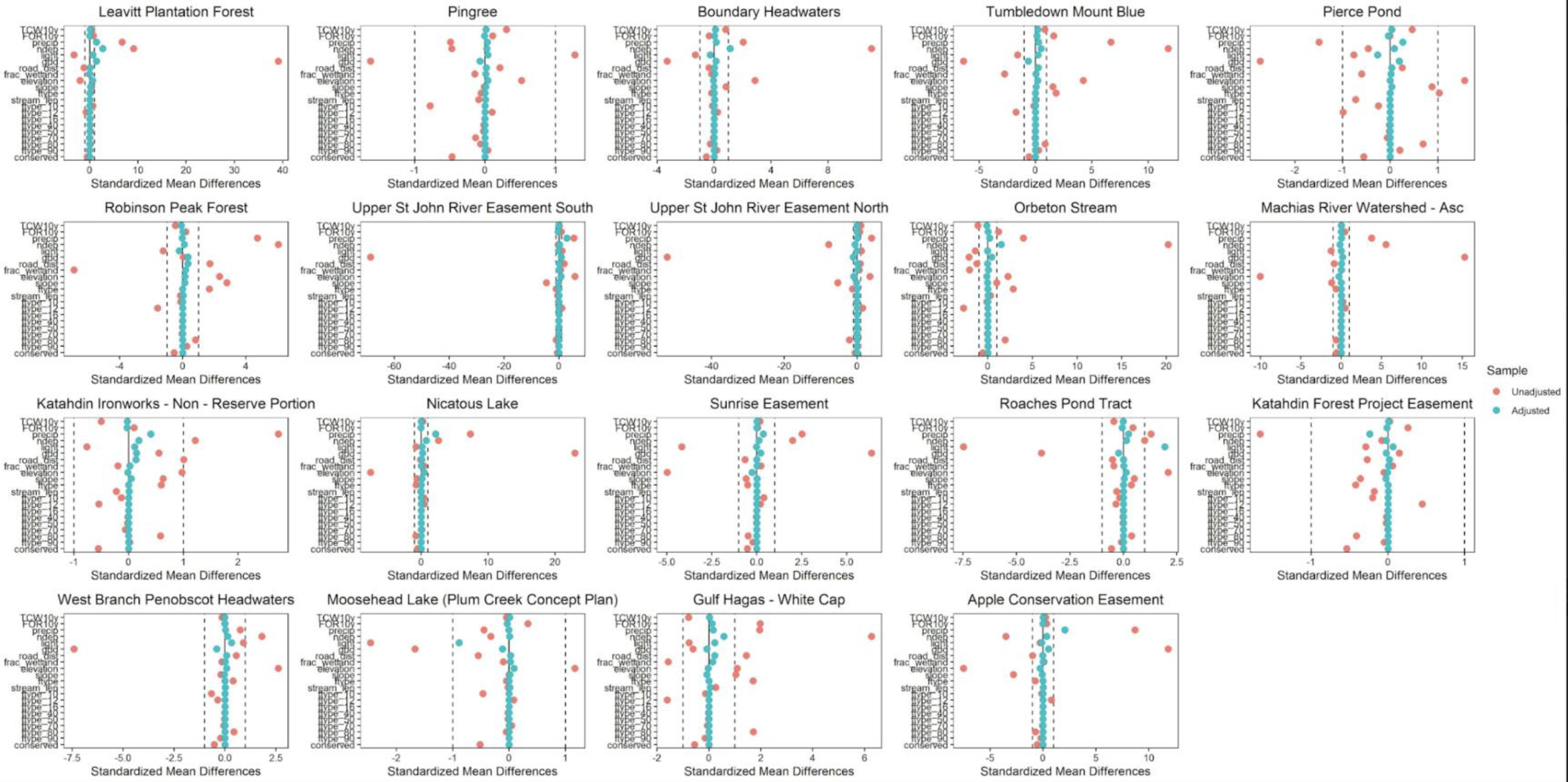
Balance Plots for covariate matching for harvest.

**Table S1:** LandTrendr parameters.

| Parameter | Value |
| --- | --- |
| maxSegments | 8 |
| spikeThreshold | 0.9 |
| vertexCountOvershoot | 3 |
| preventOneYearRecovery | true |
| recoveryThreshold | 0.75 |
| pvalThreshold | 0.05 |
| bestModelProportion | 0.75 |
| minObservationsNeeded | 6 |

**Table S2.**
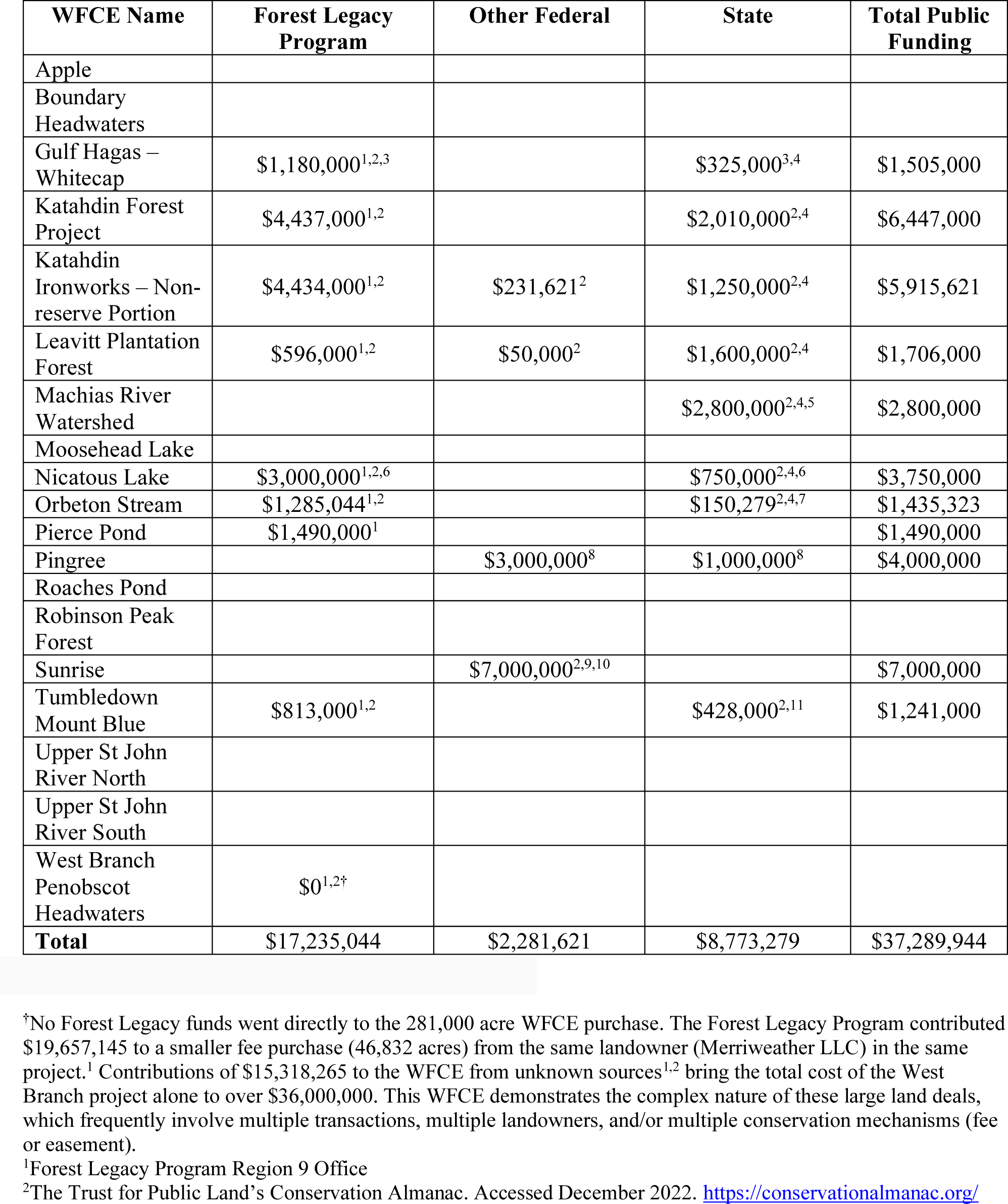
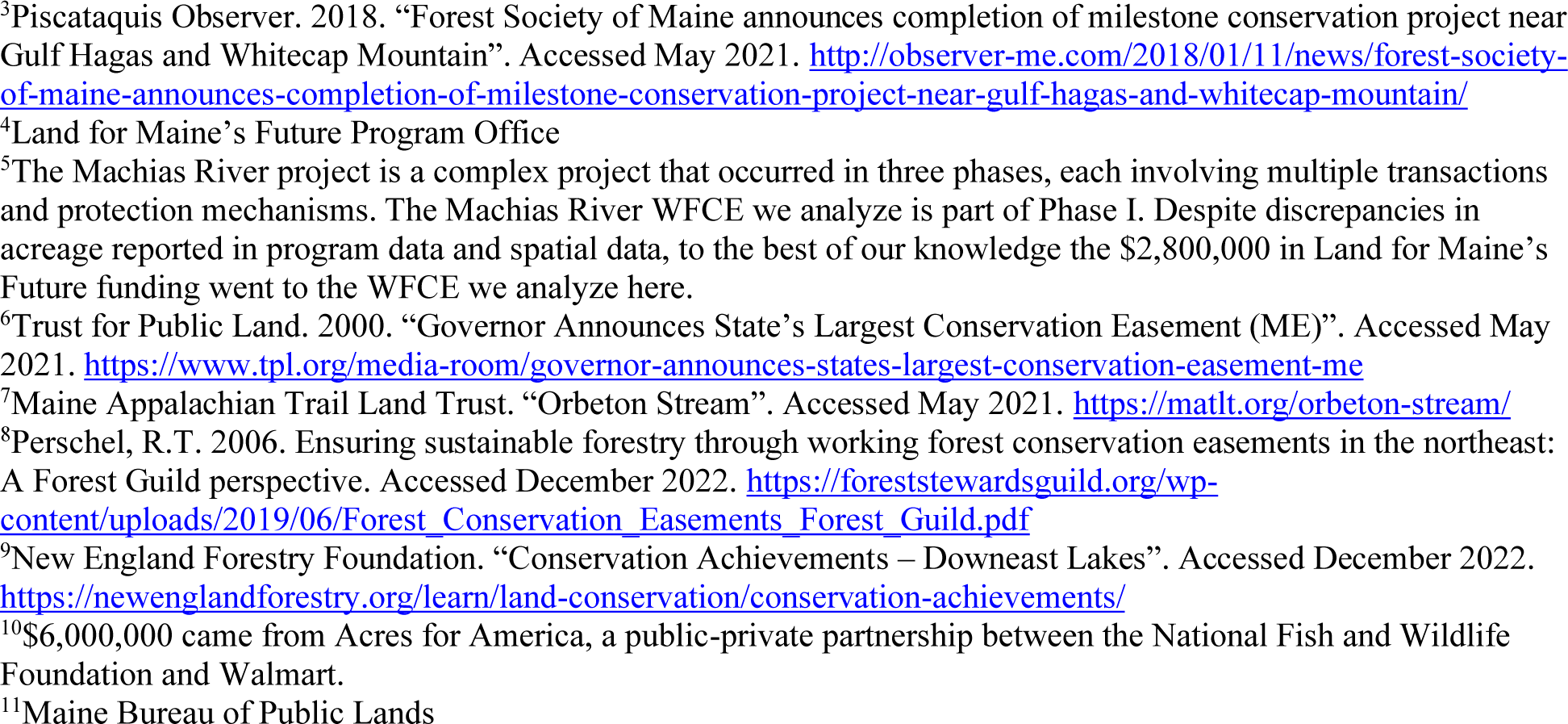
Public Funding for large WFCE in Maine.

| WFCE Name | Forest Legacy Program | Other Federal | State | Total Public Funding |
| --- | --- | --- | --- | --- |
| Apple |  |  |  |  |
| Boundary Headwaters |  |  |  |  |
| Gulf Hagas – Whitecap | \$1,180,000<sup>1,2,3</sup> | | \$325,000<sup>3,4</sup> | \$1,505,000 |
| Katahdin Forest Project | \$4,437,000<sup>1,2</sup> | | \$2,010,000<sup>2,4</sup> | \$6,447,000 |
| Katahdin Ironworks – Non-reserve Portion | \$4,434,000<sup>1,2</sup> | \$231,621<sup>2</sup> | \$1,250,000<sup>2,4</sup> | \$5,915,621 |
| Leavitt Plantation Forest | \$596,000<sup>1,2</sup> | \$50,000<sup>2</sup> | \$1,600,000<sup>2,4</sup> | \$1,706,000 |
| Machias River Watershed | | | \$2,800,000<sup>2,4,5</sup> | \$2,800,000 |
| Moosehead Lake |  |  |  |  |
| Nicatous Lake | \$3,000,000<sup>1,2,6</sup> | | \$750,000<sup>2,4,6</sup> | \$3,750,000 |
| Orbeton Stream | \$1,285,044<sup>1,2</sup> | | \$150,279<sup>2,4,7</sup> | \$1,435,323 |
| Pierce Pond | \$1,490,000<sup>1</sup> | | | \$1,490,000 |
| Pingree | | \$3,000,000<sup>8</sup> | \$1,000,000<sup>8</sup> | \$4,000,000 |
| Roaches Pond |  |  |  |  |
| Robinson Peak Forest |  |  |  |  |
| Sunrise | | \$7,000,000<sup>2,9,10</sup> | | \$7,000,000 |
| Tumbledown Mount Blue | \$813,000<sup>1,2</sup> | | \$428,000<sup>2,11</sup> | \$1,241,000 |
| Upper St John River North |  |  |  |  |
| Upper St John River South |  |  |  |  |
| West Branch Penobscot Headwaters | \$0<sup>1,2†</sup> | | | |
| **Total** | \$17,235,044 | \$2,281,621 | \$8,773,279 | \$37,289,944 |

†No Forest Legacy funds went directly to the 281,000 acre WFCE purchase. The Forest Legacy Program contributed \$19,657,145 to a smaller fee purchase (46,832 acres) from the same landowner (Merriweather LLC) in the same project.<sup>1</sup> Contributions of \$15,318,265 to the WFCE from unknown sources<sup>1,2</sup> bring the total cost of the West Branch project alone to over \$36,000,000. This WFCE demonstrates the complex nature of these large land deals, which frequently involve multiple transactions, multiple landowners, and/or multiple conservation mechanisms (fee or easement).
<sup>1</sup>Forest Legacy Program Region 9 Office
<sup>2</sup>The Trust for Public Land's Conservation Almanac. Accessed December 2022. https://conservationmanac.org/
<sup>3</sup>Piscataquis Observer. 2018. “Forest Society of Maine announces completion of milestone conservation project near Gulf Hagas and Whitecap Mountain”. Accessed May 2021. http://observer-me.com/2018/01/11/news/forest-society-of-maine-announces-completion-of-milestone-conservation-project-near-gulf-hagas-and-whitecap-mountain/
<sup>4</sup>Land for Maine’s Future Program Office
<sup>5</sup>The Machias River project is a complex project that occurred in three phases, each involving multiple transactions and protection mechanisms. The Machias River WFCE we analyze is part of Phase I. Despite discrepancies in acreage reported in program data and spatial data, to the best of our knowledge the \$2,800,000 in Land for Maine’s Future funding went to the WFCE we analyze here.
<sup>6</sup>Trust for Public Land. 2000. “Governor Announces State’s Largest Conservation Easement (ME)”. Accessed May 2021. https://www.tpl.org/media-room/governor-announces-states-largest-conservation-easement-me
<sup>7</sup>Maine Appalachian Trail Land Trust. “Orbeton Stream”. Accessed May 2021. https://matlt.org/orbeton-stream/
<sup>8</sup>Perschel, R.T. 2006. Ensuring sustainable forestry through working forest conservation easements in the northeast: A Forest Guild perspective. Accessed December 2022. https://foreststewardsguild.org/wp-content/uploads/2019/06/Forest_Conservation_Easements_Forest_Guild.pdf

**Table S3.**
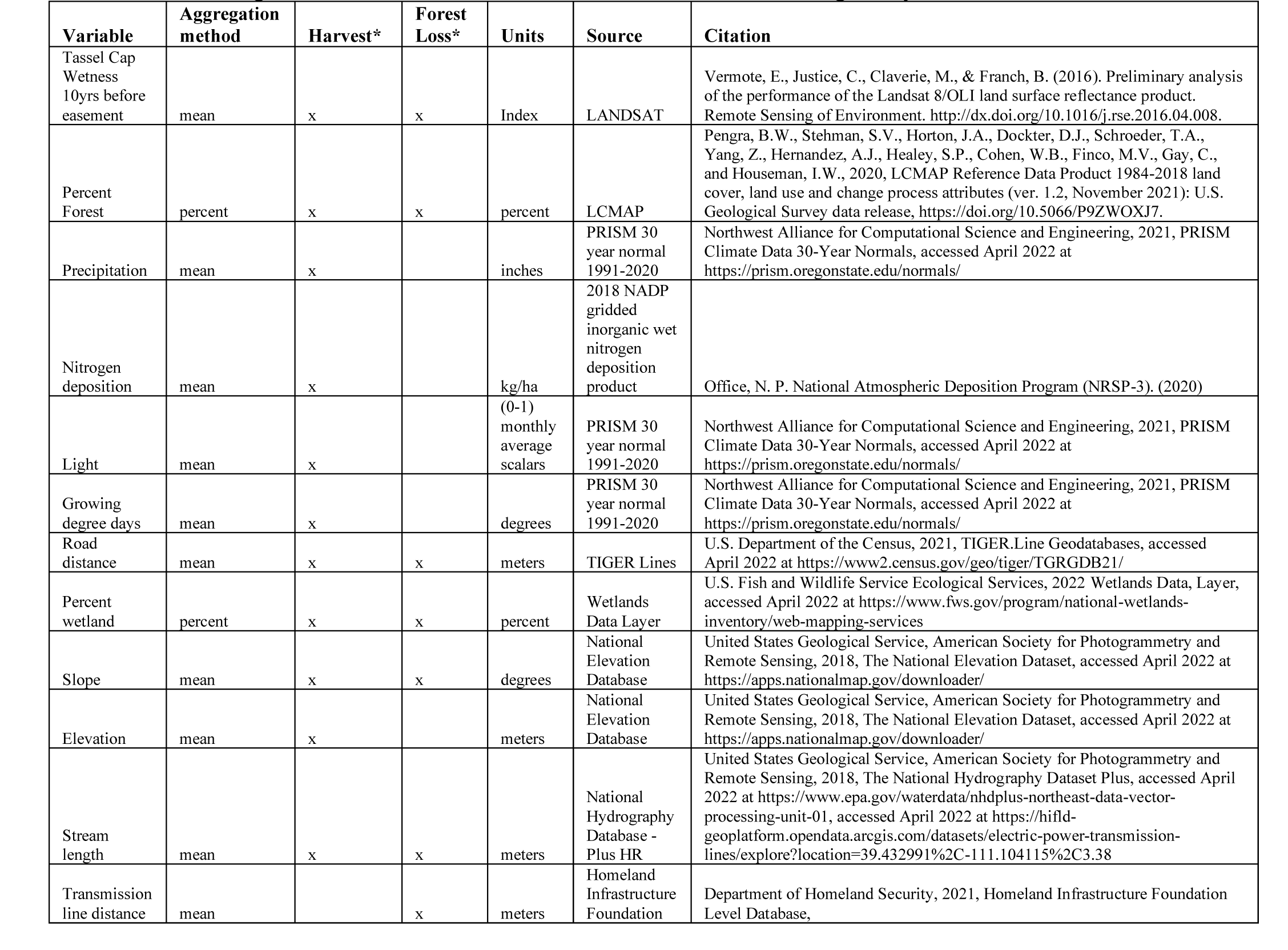
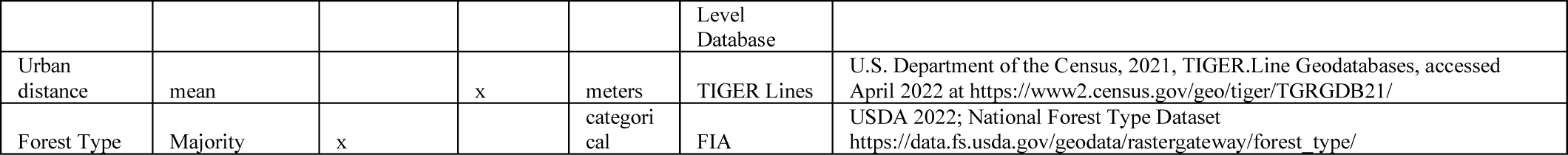
Matching variables used in Forest loss and Forest harvesting analysis.

## Footnotes

1 Please see interactive harvest viewer associated with Pasquarella et al (in review) on Google Earth Engine here: https://valeriepasquarella.users.earthengine.app/view/harvest-map-explorer

## Literature Cited

1. Bliss, J. C., Kelly, E. C., Abrams, J., Bailey, C., & Dyer, J. (2010). Disintegration of the U. S. industrial forest estate: Dynamics, trajectories, and questions. Small-scale Forestry, 9, 53–66.

2. Brown, J. F., Tollerud, H. J., Barber, C. P., Zhou, Q., Dwyer, J.L., Vogelmann, J. E., Loveland, T. R., Woodcock, C. E., Stehman, S. V., Zhu, Z., Pengra, B. W., Smith, K., Horton, J. A., Xian, G., Auch, R. F., Sohl, T. L., Sayler, K. L., Gallant, A. L., Zelenak, D., Reker, R. R., & Rover, J. (2020). Lessons learned implementing an operational continuous United States national land change monitoring capability: The Land Change Monitoring, Assessment, and Projection (LCMAP) approach. Remote Sensing of Environment, 238, 111356.

3. Congressional Research Service (CRS). (2017). Agricultural Conservation: A Guide to Programs, R40763

4. Congressional Research Service (CRS). (2018). Forest Service Assistance Programs, R45219

5. Congressional Research Service (CRS). (2019). Charitable Conservation Contributions: Potential for Abuse?

6. Daigle, J.J., Utley, L., Chase, L. C., Kuentzel, W., & Brown, T. (2012). Does new large landownership and their management priorities influence public access in the Northern Forest. Journal of Forestry, 110(2), 89–96.

7. Gunn, J. S., Ducey, M. J., & Belair, E. P. (2019). Evaluating Degradation in a North American Temperate Forest. Forest Ecology and Management, 432, 415–426.

8. Gunnoe, A., Bailey, C., & Ameyaw, L. (2018). Millions of Acres, Billions of Trees: Socioecological Impacts of Shifting Timberland Ownership. Rural Sociology, 83, 799–822.

9. Harvard Forest. (2023). New England Protected Open Space v1.2 [Dataset]. doi:10.5281/zenodo.7764284

10. Ho, D. E., King, G., Stuart, E. A., & Imai, K. (2011). MatchIt: Nonparametric Preprocessing for Preprocessing for Parametric Causal Inference. Journal of Statistical Software, 42, 1–28.

11. Kay, K. (2016). Breaking the bundle of rights: Conservation easements and the legal geographies of individuating nature. Environment and Planning A, 48(3), 504–522.

12. Kennedy, R. E., Yang, Z., Gorelick, N., Braaten, J., Cavalcante, L., Cohen, W. B., & Healey, S. (2018). Implementation of the LandTrendr algorithm on google earth engine. Remote Sensing, 10(5), 691.

13. L’Roe, A. W., & Rissman, A. R. (2017). Factors that influence working forest conservation and parcelization. Landscape and Urban Planning, 167, 14–24.

14. Legaard, K. R., Sader, S. A., & Simons-Legaard, E. M. (2015). Evaluating the impact of abrupt changes in forest policy and management practices on landscape dynamics: Analysis of a landsat image time series in the Atlantic Northern forest. PLoS ONE, 10, 1–24.

15. Lovering, J., Swain, M., Blomqvist, L., & Hernandez, R. R. (2022). Land-use intensity of electricity production and tomorrow’s energy landscape. PLoS ONE, 17, 1–17.

16. Merrow, M. (2018). Towers, Trees, and Transmission Lines: The Fight between Property Rights, Power, and Profit. Hastings Environmental Law Journal, 24.

17. Meyer, S. R., Cronan, C. S., Lilieholm, R. J., Johnson, M. L., & Foster, D. R. (2014). Land conservation in northern New England: Historic trends and alternative conservation futures. Biological Conservation, 174, 152–160.

18. Murray, H., Catanzaro, P., Markowski-Lindsay, M., Butler, B., & Eichman, H. (2018). Economic Contributions of Land Conserved by the USDA Forest Service’s Forest Legacy Program. Forest Science, 67, 629–632.

19. Nolte, C., Meyer, S. R., Sims, K. R. E., & Thompson, J. R. (2019). Voluntary, permanent land protection reduces forest loss and development in a rural-urban landscape. Conservation Letters, 33, 1035–1044.

20. Noone, M. D., Sader, S. A., & Legaard, K. R. (2012). Are forest disturbance rates and composition influenced by changing ownerships, conservation easements, and land certification? Forest Science, 58, 119–129.

21. Owley, J., & Rissman, A. R. (2016). Trends in private land conservation: Increasing complexity, shifting conservation purposes and allowable private land uses. Land Use Policy, 51, 76–84.

22. Parker, D. P., & Thurman, W. N. (2019). Private Land Conservation and Public Policy: Land Trusts, Land Owners, and Conservation Easements. Annual Review of Resource Economics, 11, 337–354.

23. Pasquarella, V. J., Morreale, L. L., Brown, C., Kilbride, J. B., & Thompson, J. R. (In Review). Not-so-random forests: Comparing voting and decision tree ensembles for characterizing partial harvest events in complex forested landscapes. International Journal of Applied Earth Observation and Geoinformation, in review.

24. Pidot, J. (2005). Reinventing Conservation Easements: A Critical Examination and Ideas for Reform. Page Lincoln Institute of Land Policy Working Paper.

25. Reeves, T., Mei, B., Siry, J., Bettinger, P., & Ferreira, S. (2020). Towards a characterization of working forest conservation easements in Georgia, USA. Forests, 11, 1–17.

26. Rissman, A., Bihari, M., Hamilton, C., Locke, C., Lowenstein, D., Motew, M., Price, J., & Smail, R. (2013). Land management restrictions and options for change in perpetual conservation easements. Environmental Management, 52, 277–288.

27. Sass, E. M., Butler, B. J., & Markowski-Lindsay, M. A. (2020). Forest ownership in the conterminous United States circa 2017: distribution of eight ownership types - geospatial dataset. Fort Collins, CO.

28. Sass, E. M., Markowski-Lindsay, M., Butler, B. J., Caputo, J., Hartsell, A., Huff, E., & Robillard, A. (2021). Dynamics of Large Corporate Forestland Ownerships in the United States. Journal of Forestry, 1–13.

29. Saul, R. S. (2021). The nature of financialized nature: The future of institutional investments in U.S. food and fiber production in the climate emergency era. Harvard Forest Research Paper 36. Petersham, MA.

30. Sayen J. (2024). Children of the Northern Forest; Wild New England’s History from Glaciers to Global Warming. Yale University Press.

31. Sims K. E., Thompson J.R., Meyer S., Nolte C., Plisinski, J. (2019) Assessing the local economic impacts of land protection. Conservation Biology. 33(5) 1035–1044

32. Stein, P. R. (2011). Trends in forestland ownership and conservation. Forest History Today, 83–86.

33. Tesini, D. (2009). Working Forest Conservation Easements. The Urban Lawyer, 41, 359–375.

34. Thompson, J. R., Canham, C., Morreale, L., Kittredge, D. B., & Butler, B. J. (2017a). Social and Biophysical Variation in Regional Timber Harvest Regimes. Ecological Applications, 27(3), 942–955.

35. Thompson, J. R., Carpenter, D. N., Cogbill, C. V., & Foster, D. R. (2013). Four Centuries of Change in Northeastern United States Forests. PLoS ONE, 8, 1–15.

36. Thompson, J. R., Plisinski, J., Olofsson, P., Holden, C. E., & Duveneck, M. J. (2017b). Forest loss in New England: A projection of recent trends. PLoS ONE, 12, 1–17.

37. Thompson, J. R., Plisinski, J. S., Lambert, K. F., Duveneck, M. J., Morreale, L., McBride, M., MacLean, M. G., Weiss, M., & Lee, L. (2020). Spatial Simulation of Codesigned Land Cover Change Scenarios in New England: Alternative Futures and Their Consequences for Conservation Priorities. Earth’s Future, 8.

38. USFS (2017) Forest Legacy Program Implementation Guidelines. U.S. Forest Service. FS 1088. Washington D.C.

39. Zhao, J., Daigneault, A., & Weiskittel, A. (2020). Forest landowner harvest decisions in a new era of conservation stewardship and changing markets in Maine, USA. Forest Policy and Economics, 118, 102251.

40. Zhu, Z., & Woodcock, C. E. (2014). Continuous change detection and classification of land cover using all available Landsat data. Remote Sensing of Environment, 144, 152–171.

## Supplemental References not in main Literature Cited section

1. Kennedy, R. E., Yang, Z., & Cohen, W. B., (2010). Detecting trends in forest disturbance and recovery using yearly Landsat time series: 1. LandTrendr – Temporal segmentation algorithms. Remote Sensing of Environment, 114, 2897–2910.

2. Loeb, C.D., & D’Amato, A.W. (2019). “New Hybrid Protected Lands Layer for Vermont Conservation Design Analysis (February 2019)” [Dataset]. Rubenstein School Masters Project Publications. 23. https://scholarworks.uvm.edu/rsmpp/23

3. Pasquarella, V. J., Arévalo, P., Bratley, K. H., Bullock, E. L., Gorelick, N., Yang, Z., & Kennedy, R. E. (in review). Demystifying LandTrendr and CCDC temporal segmentation. International Journal of Applied Earth Observation and Geoinformation, 110, 102806.

